# RNA Editing Signatures Predict Response to Immunotherapies in Melanoma Patients

**DOI:** 10.1101/2020.08.12.248393

**Authors:** Jalal Siddiqui, Wayne O. Miles

## Abstract

Immunotherapy has improved the prognosis for half of the melanoma patients, prompting a need to understand differences between responding and non-responding patients. Gene expression profiling of tumors has focused on deriving primarily immune-related signatures, however these have shown limited predictive power. Recent studies have highlighted the role of RNA editing in modulating resistance to immunotherapy. This has led us to test whether RNA editing activity can be predictive of response in publicly available datasets of immunotherapy-treated melanoma patients. Here, we identified RNA editing signatures that were able to predict with very high accuracy and confidence patient responses and outcomes. Our analysis, however, demonstrates that RNA editing by itself is sufficient as a strong predictive tool for examining sensitivity of melanoma patients to immunotherapy.

## Main

Melanoma is a highly aggressive and frequently lethal cancer. The advent of immunotherapy has significantly improved the prognosis for around half of melanoma patients, however our understanding of which patients will and will not respond to these treatments is a significant hurdle^1–4^. A number of groups have profiled the tumors of melanoma patients on immunotherapy clinical trials utilizing RNA-sequencing (RNA-Seq) in an attempt to identify gene expression signatures associated with patient response^2–4^. Each of these studies have provided important biological insights into genomic and transcriptional changes that drive melanoma, however the predictive power of these signatures is limited.

Recent studies have highlighted the role of RNA editing, particularly by the adenosine deaminase acting on RNA (ADAR) family of proteins as important for the immune response and T-cell activation^5–9^. The ADAR protein family (ADAR1-3 (ADAR3: is enzymatically inactive)) catalyzes the deamination of Adenines (A) within doublestranded regions of RNA (dsRNA) into Inosines (I), in a process known as A-to-I editing^10^. The resulting I-U base-pair is significantly less stable than the replaced A-U base-pair causing destabilization of dsRNA^10^. These modifications have vital roles in cellular homeostasis, as unedited dsRNA regions are recognized by human cells as viral contaminants and trigger strong immune responses^7–10^. Importantly for our study, the resulting I base is recognized as a Guanine (G) during library construction, enabling the identification of A-I editing sites from RNA-Seq data using a number of available computational tools^11–13^. Based on the strong link between ADAR activity, the immune response and a real need to determine immunotherapy response in patients, we tested the predictive power of RNA editing in publicly available datasets^2,3,11^.

## Results

Utilizing the Hugo dataset^2^, we first examined whether the levels of interferon response or ADAR genes correlated with patient response to immunotherapy. From these results, we found that the expression of interferon (Figure 1a) or ADAR genes (Figure 1b) is not associated with patient outcome (Non-Responder (NR) vs. Responder (R)). In addition, the overall levels of A-I editing events, as measured by AG/TC transitions, also appeared random (Figure 1c, Table S1). We next asked whether RNA editing sites (RES) in independent genes were associated with patient response. For this, we tabulated the RES score^14^ for each gene and identified gene RES scores significantly associated with response (t-test p<0.05) and not differentially expressed (Wald p>0.05)^15–16^. From this analysis, we identified 13 up-regulated (Figure 1d) and 248 down-regulated RES scores (Figure 1e, Table S2) that correlated with patient response to immunotherapy. Down-regulated RES events provided the cleanest clustering of patients based on outcome (Figure 1e). To determine the heterogeneity between patients, we compared the means of the signature RES scores and found a striking and statistically significant difference between non-responding vs responding patients (Figure 1f, Table S3). These findings suggest that RES scores may provide the basis for more accurately predicting patient response to immunotherapy.

**Figure 1:**
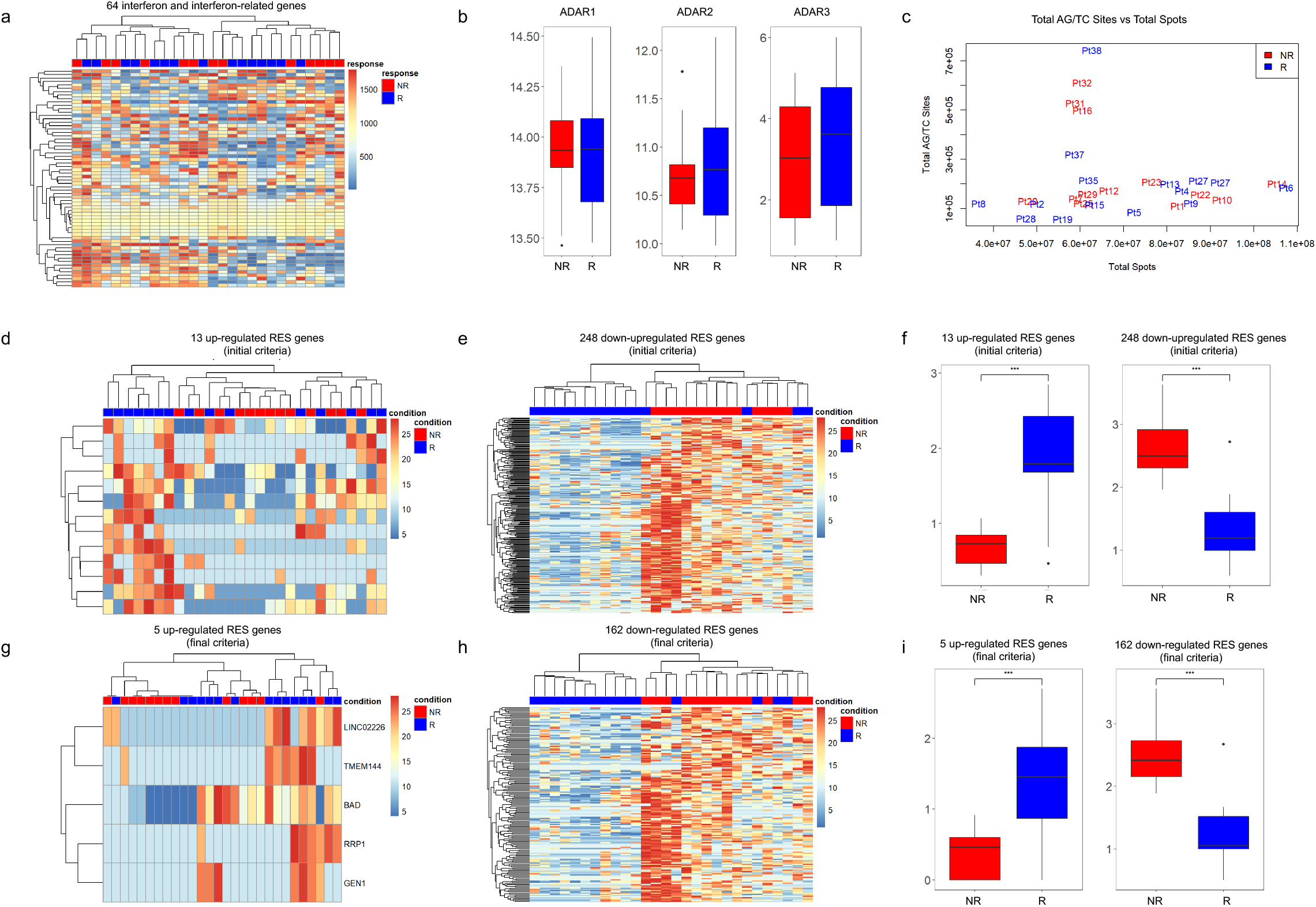
RNA editing sites can segregate melanoma patients based on response to immunotherapy. **a)** Heat map of Interferon and Interferon-related gene expression changes in non-responder (NR) and responder (R) patients from the Hugo dataset. **b)** ADAR1, ADAR2 and ADAR3 gene expression levels in non-responder and responder patients. **c)** Total AG/TC RNA editing sites vs. total spots in non-responder (red) and responder (blue) patients. **d)** Heat map of up-regulated RES scores in genes using initial criteria in non-responder and responder patients. **e)** Heat map of down-regulated RES scores in genes using initial criteria in non-responder and responder patients. **f)** Means of RES scores of up-regulated and down-regulated genes using initial criteria in nonresponder and responder patients. **g)** Heat map of up-regulated RES scores in genes using final criteria in non-responder and responder patients. **h)** Heat map of down-regulated RES scores in genes using final criteria in non-responder and responder patients. **i)** Means of RES scores of up-regulated and down-regulated genes using final criteria in non-responder and responder patients. *p<0.05, **p<0.01, ***p<0.001

Analysis of this type can be affected by confounding factors. To minimize this, we added two additional filtering steps. First, we removed RES scores that had significant linear relationships (p>0.05) to transcript levels. Second, we used linear regression on the RES scores with the predictors being response and transcript levels. Following this filtering, we retained only RES scores with a significant response coefficient (p<0.05) and removed the confounding effect of transcriptional levels. This approach enabled the identification of 5 up-regulated and 162 down-regulated RES scores that can classify responding vs. non-responding patients (Figures 1g and 1h, Table S2). In agreement with our findings based on the initial criteria, we note that down-regulated RES scores present the most accurate patient clustering (Figure 1h). The mean RES scores for each of these groups was again tightly correlated with outcome (Figure 1i, Table S3). Importantly, the mean gene expression of these genes was unchanged in responders vs. non-responders (Figure S1a, 1b, Table S3). Collectively, these results show that RES signatures can separate patients by outcome to immunotherapy and that these changes are driven by RNA editing activity and not transcription.

The predictive power of the RES score in the Hugo dataset, compelled us to investigate whether RNA editing signatures could stratify patients from additional cohorts. For this, we selected the Riaz melanoma dataset that included RNA-Seq for melanoma patients treated with nivolumab from the following groups: responder, non-responder, and stable disease patients^3^. In agreement with the Hugo data, an interferon response signature (Figure 2a), ADAR levels (Figure 2b) and gross A-I editing sites (Figure 2c, Table S4), were not predictive. Utilizing our initial RES criteria, we identified gene RES scores that correlated with patient outcome (Figure 2d, 2e, Table S5). This is particularly striking for down-regulated RES sites, as patients that respond to immunotherapy cluster very tightly together (Figure 2e). Mean RES scores across these patient groups also show strong separation in responding patients (Figure 2f, Table S6). Interestingly, patients with stable disease frequently correlate with RES scores of non-responders and show similar RES means (Figure 2d-f). As above, we then filtered these events to remove confounding effects and identified up-regulated (Figure 2g) and down-regulated (Figure 2h) RES scores that tightly correlated with patient outcome. The mean RES scores for each responder is highly significant (Figure 2i, Table S6) and are unchanged at the transcript level (Figure S1c, 1d, Table S6). Collectively, these findings confirm that RES score analysis can be utilized to segregate patients based on their response to immunotherapy. This approach can be used on diverse datasets with differences in sequencing coverage, patient population and immunotherapy agent.

**Figure 2:**
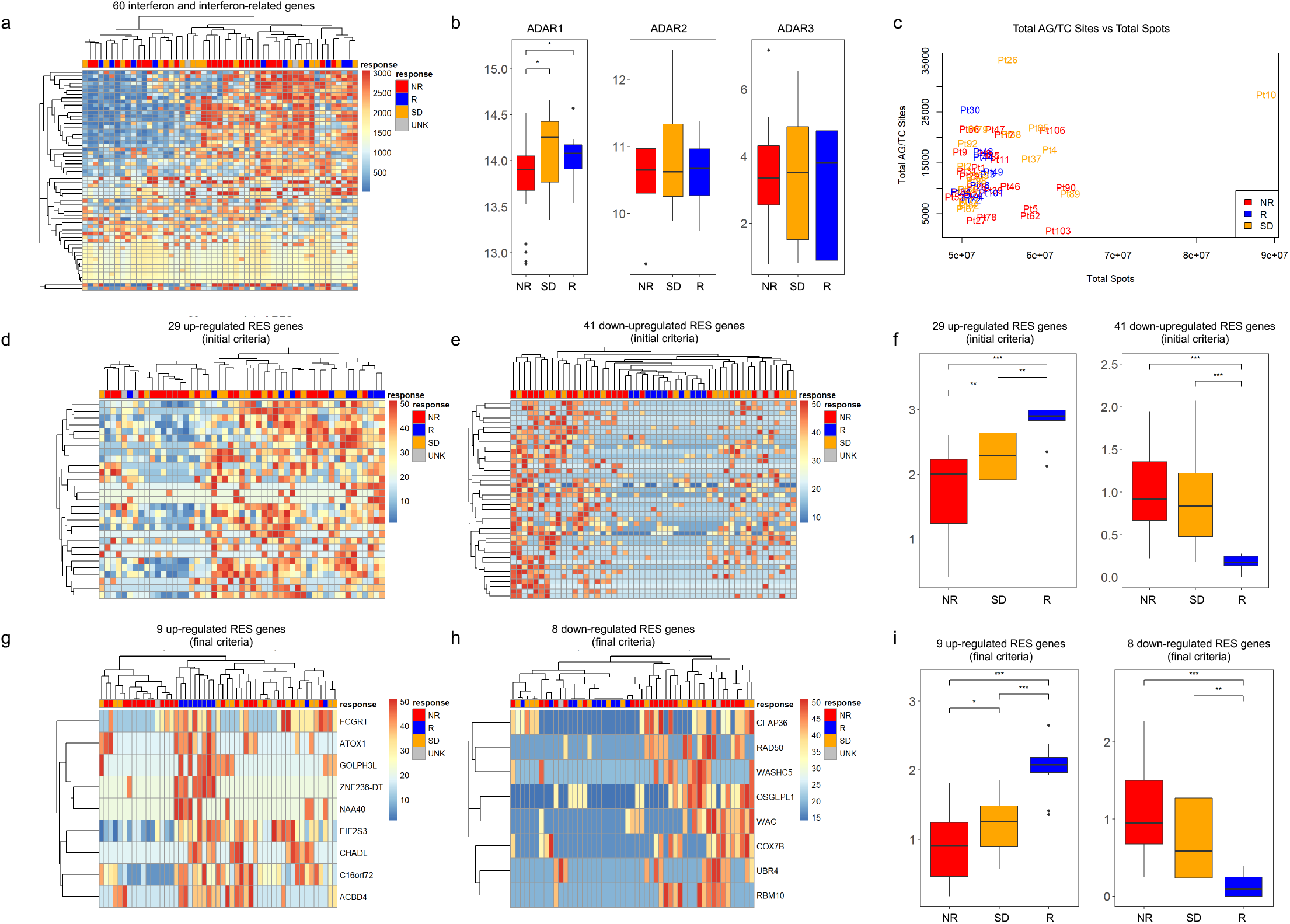
RES scores can be used across datasets to sub classify immunotherapy response. **a)** Heat map of Interferon and Interferon-related gene expression changes in non-responder (NR), responder (R), stable disease (SB) and unknown (UNK) patients from the Riaz dataset. **b)** ADAR1, ADAR2 and ADAR3 gene expression levels in nonresponder, stable disease and responder patients. **c)** Total AG/TC RNA editing sites vs. total spots in non-responder (red), responder (blue) and stable disease (orange) patients. **d)** Heat map of up-regulated RES scores in genes using initial criteria in non-responder, responder, stable disease and unknown patients. **e)** Heat map of down-regulated RES scores in genes using initial criteria in non-responder, responder, stable disease and unknown patients. **f)** Means of RES scores of up-regulated and down-regulated genes using initial criteria in non-responder, stable disease and responder patients. **g)** Heat map of up-regulated RES scores in genes using final criteria in non-responder, responder, stable disease and unknown patients. **h)** Heat map of down-regulated RES scores in genes using final criteria in non-responder, responder, stable disease and unknown patients. **i)** Means of RES scores of up-regulated and down-regulated genes using final criteria in non-responder, stable disease and responder patients. *p<0.05, **p<0.01, ***p<0.001

Our RES scores outlined in Figure 1 and 2 correlate with patient response, however we wanted to investigate the predictive power of this pipeline for patients. For this, we used logistic regression models from up and down-regulated mean RES scores from each dataset. Using the Hugo dataset, we found that up-regulated RES scores had a greater than 78% capacity to predict patients that will respond to immunotherapy (Figure 3a, S2a-c, Table S7). We note that the sole responding patient that our model does not predict (blue dot below 0.5 in Figure 3a), is patient 38 (Pt38), which had very high AG/TC levels and is an outlier in Figure 1c. In contrast, the down-regulated RES scores have an over 96% predictive capacity and separates all of the responding patients (Figure 3b, S2d-f, Table S7). In the independent Riaz dataset, we also find that our logistic regression of RES scores predicts patient response with both the up (90%) (Figure 3c, S3a-c) and down-regulated RES genes (Figure 3d, S3d-f, Table S7) having a greater than 87% predictive capacity. Based on these analyses, the down-regulated RES scores represent the most accurate model for determining patient sensitivity. These results highlight the predictive power of this approach for identifying patients that are likely to respond to immunotherapy.

**Figure 3:**
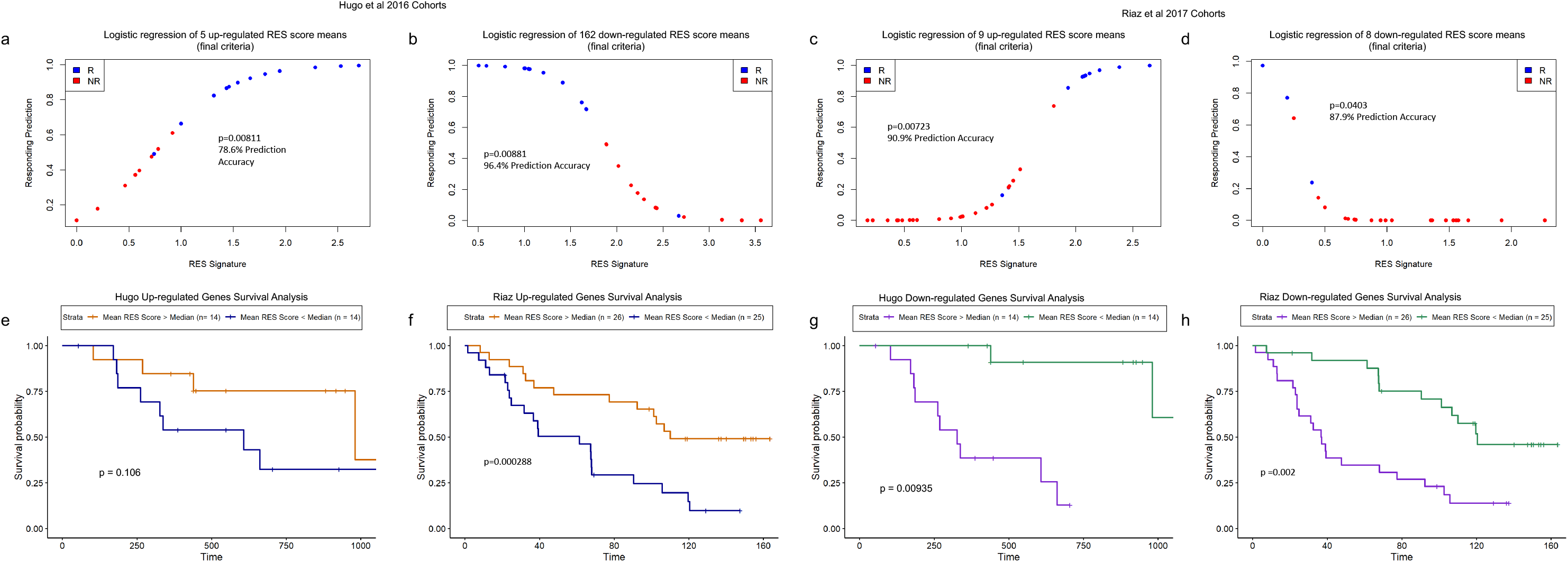
RES scores accurately predict response and survival of melanoma patients to immunotherapy. **a)** Logistic regression models from the Hugo dataset for up-regulated RES score means and responding prediction for responder (R, blue) and non-responder (NR, red) patients. **b)** Logistic regression models from the Hugo dataset for down-regulated RES score means and responding prediction for responder and nonresponder patients. **c)** Logistic regression models from the Riaz dataset for up-regulated RES score means and responding prediction for responder and non-responder patients. **d)** Logistic regression models from the Riaz dataset for down-regulated RES score means and responding prediction for responder and non-responder patients. **e)** Survival analysis of patients stratified by upper 50% (red) and lower 50% (blue) means of up-regulated RES scores for genes in the Hugo cohort. **f)** Survival analysis of patients stratified by upper 50% (red) and lower 50% (blue) means of up-regulated RES scores for genes in the Riaz cohort. **g)** Survival analysis of patients stratified by upper 50% (purple) and lower 50% (green) means of down-regulated RES scores for genes in the Hugo cohort. **h)** Survival analysis of patients stratified by upper 50% (purple) and lower 50% (green) means of down-regulated RES scores for genes in the Riaz cohort.

We next evaluated how these RES scores correlated with patient survival. For this, patients were stratified by mean RES scores. For patients with elevated RES scores in the up-regulated group, we find improved survival periods for patients within the Hugo cohort (Figure 3e, Table S8). In agreement, patients with elevated RES score means within the Riaz dataset, show significant survival differences to those with lower means (Figure 3f, Table S8). Although these findings strongly support the predictive power of our model, the down-regulated RES score means show the most significant differences in patient survival (Figure 3g-h, Table S8). In this data, patients with lower mean RES scores display real survival benefit that is independent of the dataset. We next tested how this approach would cluster stable disease patients, and found that this logistic regression test was highly accurate and would cluster these patients with non-responders (Figure S4). Based on the logistic regression model and survival analysis of these patients, the down-regulated RES score enables the most accurate sub-classification of melanoma patients across datasets. This data highlights the predictive nature of RES scores for understanding the clinical benefit for patients on immunotherapy.

We next examined the predictive power of recurrent RES sites within different patient groups. For this, we sub-classified RES sites enriched from responder or non-responder patients. Using recurrent RES sites, we were able to completely separate patients within the Hugo cohort based on clinical response (Figure 4a). This is also true for patients within the Riaz group (Figure 4b), suggesting that recurrent RNA editing within genes is contributing to response. To build on this, we compared the common genes with RES scores within both datasets. These genes with conserved A-I editing were also strongly predictive of patient response and able to segregate responders from non-responders or stable disease in each dataset (Figure 4c, 4d). We then examined whether recurrent RES sites existed between datasets that could predict patient response. For this analysis, we compared RES sites from both datasets and identified 5 RES sites that were present and significant in both (Table S9). We next tested whether these recurrent RES events could separate patients by response, and found that although the number of RES sites is small, these sites correlated with patient response (Figure 4e, 4f). By using the cumulative levels of RES sites within these genes, we find that elevated numbers of RES sites correlate with improved patient survival in both datasets (Figure 4g, 4h). To evaluate the hazard potential of this metric, we utilized Cox Proportional Hazards modeling^17^. From this, we find significant associations between patient response and the number of common RES sites (Table S3, Table S9). Our findings strongly implicate recurrent RES sites within melanoma as predictive of patient outcome and survival.

**Figure 4:**
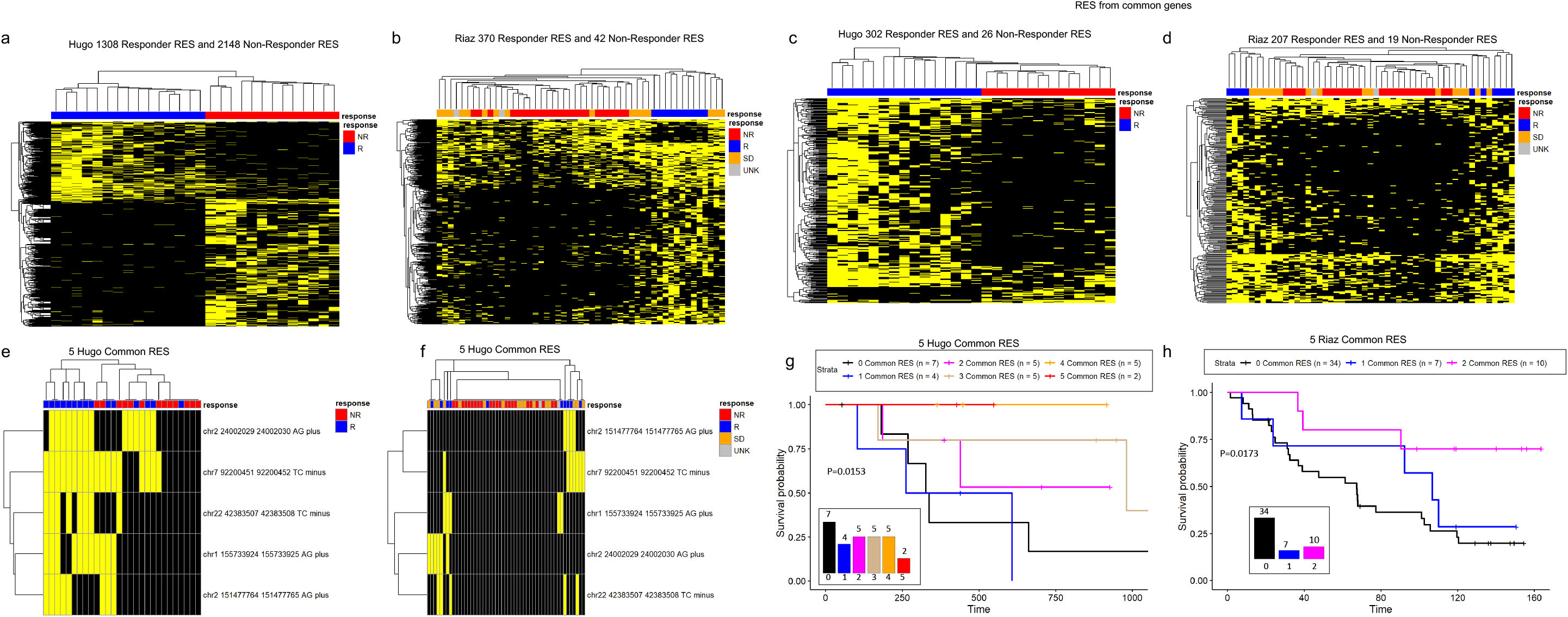
Recurrent and common RES sites occur and predict outcomes in melanoma tumors. **a)** Heat map of significant recurrent RES sites in non-responder (NR) and responder (R) patients from the Hugo dataset. **b)** Heat map of significant recurrent RES sites in non-responder (NR), responder (R), stable disease (SD) and unknown (UNK) patients from the Riaz dataset. **c)** Heat map of significant shared RES genes in non-responder and responder patients from the Hugo dataset. **d)** Heat map of significant shared RES genes in non-responder, responder, stable disease and unknown patients from the Riaz dataset. **e)** Heat map of significant shared RES sites in non-responder and responder patients from the Hugo dataset. **f)** Heat map of significant shared RES sites in responder, non-responder, stable disease and unknown patients from the Riaz dataset. **g)** Survival analysis of patients stratified by common RES number 0-5 from the Hugo dataset (0 common RES - black, 1 common RES – blue, 2 common RES – magenta, 3 common RES – tan, 4 common RES – orange, 5 common RES - red). **h)** Survival analysis of patients stratified by common RES number 0-2 from the Riaz cohort (0 common RES - black, 1 common RES – blue, 2 common RES - magenta).

## Discussion

This analysis highlights the real prognostic power of RNA editing sites and the affected genes in predicting immunotherapy responses in melanoma patients. Previous studies have focused on the analysis of genomic and transcriptomic datasets to identify immune related signatures in pathways such as interferon signaling^18^, MHC antigen presentation^3,19,20^, and innate anti-PD1 resistance^2^. However, although innovative the predictive power of these signatures to identify patient response to immunotherapy has been limited. In this study, we investigated the capacity of RNA A-I editing events to accurately predict patient outcomes and response to different immunotherapy agents. Our approach has incorporated new knowledge that has been generated by a number of groups that have identified the role of ADAR-mediated RNA editing in development and cancer^5–7,10,21,22^. These studies have linked ADAR activity or ADAR-loss to immune response levels and immune checkpoint blockade. In agreement with a number of other studies, we find no correlation with interferon signatures, ADAR levels or total A-I editing events to immunotherapy response^18,23,24^. However, by systematically identifying A-I RNA changes that do not correlate with transcriptional changes, our RES scores accurately capture A-I editing events that can predict patient response with very high levels of confidence.

These findings show that the RES score is sufficient to predict patient response on vastly different datasets. We note that the significant differences in patient population, library construction and sequencing protocols resulted in more exhaustive coverage in the Hugo dataset compared with Riaz^2,3^. In addition, patients within each trial were treated with different immunotherapies: Patients from the Hugo cohort primarily received pembrolizumab, while the Riaz cohort received nivolumab. In spite of these differences, our data suggests that the controlled use of RNA A-I editing sites is a strong predictive tool for examining the sensitivity of melanoma patients to immunotherapy.

## Data Availability

All data are presented in this manuscript are available from the corresponding author upon reasonable request.

## Acknowledgements

We would like to acknowledge the support of the Damon Runyon Cancer Research Foundation award 51-18 to W.O.M and the support of a P30 award (CA016058) to the OSUCCC from the National Cancer Institute, Bethesda, MD.

## Author Contributions

J.S. conducted the analysis and co-wrote the manuscript. W.O.M conceived the project and co-wrote the manuscript.

## Methods

### Datasets Used

The Hugo dataset (GSE78220) consisted of samples from patients treated primarily with pembrolizumab^1^. 15 were responders and 13 were non-responders based on Immune-related Response Evaluation Criteria In Solid Tumors (irRECIST) criteria^1^. The Riaz dataset (GSE91061) consisted of nivolumab-treated patients of which 23 were nonresponders, 10 responders, and 16 stable disease by RECIST criteria^2^. 2 were of unknown response designation^2^.

### RNA Editing Sites Pipeline

RNA-Seq files were imported from Gene Expression Omnibus (Hugo GSE78220 and Riaz GSE91061) using SRAToolkit 2.9.0. TrimGalore 0.6.0 was used to perform adaptor and quality trimming of RNASeq reads with the additional specification of 6 bases removed from the 5’ of each reads^3^. Bowtie2 was used for removing contaminating rRNA and tRNA reads^4^. STAR 2.5.2a was used for aligning the reads to the GRCh38 p12 genome release 31 and obtaining gene counts and BAM alignment files^5,6^. The gene counts were imported into the R environment using *DESeq2*, a Bioconductor package for differential expression analysis and the resulting data was log-regularized^7^. Additionally, a differential expression analysis was conducted between responders and nonresponders.

Sprint was used on the resulting BAM alignment files to identify RNA editing sites *de novo*.^8^ The *changesammapq.py* script from Sprint was used to convert the BAM files to the correct format for Sprint^8^. The *sprint_from_bam.py* script was used to identify regular RNA editing sites from the resulting BAM file using the GRCh8 p12 genome release 31^6^ and Sprint-provided hg38 repeat annotations. Annotating of the resulting RNA editing sites positions for genes, genomic regions, and repeats was done using functions and hg38 annotations from *annotatr*, a Bioconductor package for investigating intersecting genomic annotations^9^. Sprint-provided hg38 repeat annotations were also used in annotating the RNA editing sites.

### Identifying Differential RNA Editing

For each gene, we developed an RNA editing score defined as the number of RNA editing sites per gene^10^. For our RNA editing scores, we only focused on A to G or T to C transitions for a gene. The RNA editing scores were log2-transformed with a pseudocount of 1 to normalize the data. A two-tailed t-test was performed to determine differential RNA editing scores for each gene between responding and non-responding patients with significance criteria being two-tailed p-value < 0.05 and log2 fold change > 0.3785. Additionally, we overlapped the significant genes with DESeq2 results and removed genes that were differentially expressed at a nominal level of significance (Wald’s Test two-tailed p-values < 0.05).

A *final set* of filtering criteria (as opposed to *initial criteria* of filtering by only t-test and *DESeq* results) was developed to remove the influence of log-regularized transcript levels on number of RES identified using linear models analyzed with the *lm()* function in R. Genes with a significant *transcript* two-tailed p-value < 0.05 in *RES ~ transcript* linear model were filtered out. Additionally, the linear model *RES ~ response + transcript* was used to remove transcript levels as a confounding variable and only genes with a twotailed p-value < 0.05 for the *response* coefficient were retained.

### RNA Editing Signatures

The mean RES score of the signature up-regulated and down-regulated gene (for both initial and final criteria) was used as a measure of the RNA editing signatures for each patient sample. Two-tailed pairwise t-tests were conducted between responders, non-responders, and stable disease patients to determine how significantly each signature discriminated between response groups. Additionally, a t-test was also conducted on the corresponding log-regularized transcript levels to determine how the transcript levels of the signature genes significantly discriminated between patient response groups.

### Logistic Regression Models

The means of the significant log2-tranformed RNA editing scores from *initial* and *final* up-regulated and down-regulated genes were used as input to a set of logistic regression models using the *glm()* functions in R where *response ~ mean RNA editing score.* Twotailed p-values of the *mean RNA editing score* coefficient was used to assess significance and model accuracy was calculated for correctly classified patient samples. The *pROC* R package was used for receiver operating characteristic (ROC) analysis^11^.

### Recurrent RNA Editing Sites

AG/TC RNA editing sites from all samples in each cohort were stratified into sites identified only responders, sites identified only non-responders, and those identified in both responders and non-responders (Stable patients in Riaz were not considered). A two-tailed Fisher’s Exact test was performed to determine the contingency of RNA editing sites being enriched in responders or non-responders. Significant recurrent RNA editing sites had a two-tailed p-value < 0.05 and were only annotated to genes with DESeq2 nominal Wald’s Test p-values > 0.05. Significant sites whose contingency odds ratio favored responders were responder-enriched and significant sites whose odds ratios favored non-responders were non-responder enriched. For the responder and nonresponder enriched RES we determined the annotated genes for these sites using functions derived from *annotatr*^9^. The responder and non-responder enriched RES and their annotated genes were compared across the Hugo and Riaz cohorts.

### Survival Analysis

Survival analyses were done to determine the effects of RNA editing signatures and recurrent RNA editing sites on patient survival. Analyses were performed on all patient samples within a cohort. The *survival* and *survminer* R packages were used^12^. A survival object or response variable was created from survival data using the *Surv()* function. Kaplan-Meir curves were created using the *survfit()* function and Cox proportional hazard regression modeling was done via the *coxph()* function^13^. The *ggsurvplot()* function was used for visualizing survival curves.

---

**Figure S1:**
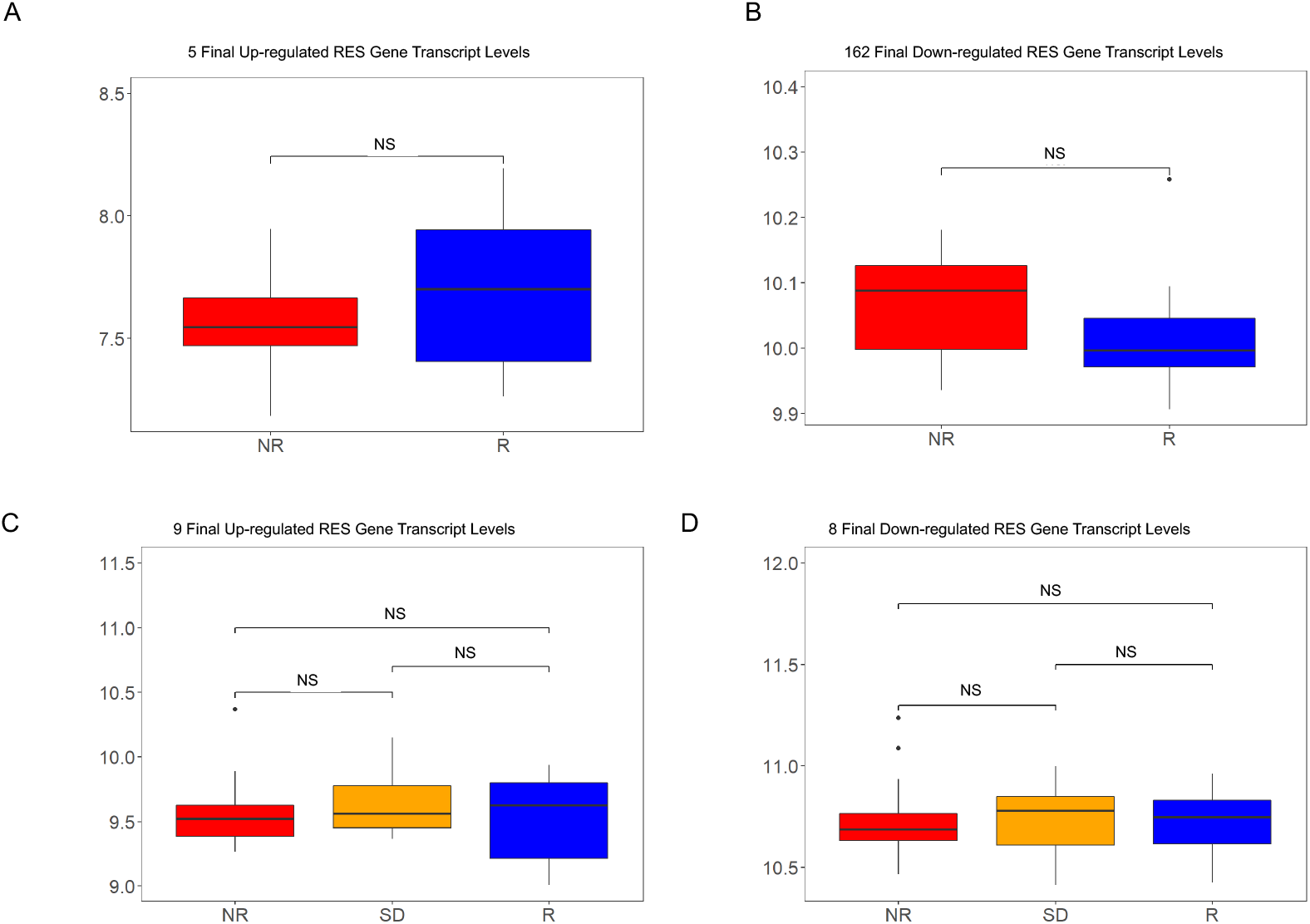
Gene expression levels of RES containing genes do not correlate with outcome. **a)** Expression levels of predictive genes that contain up-regulated RES scores in the Hugo dataset. **b)** Expression levels of predictive genes that contain down-regulated RES scores in the Hugo dataset. **c)** Expression levels of predictive genes that contain up-regulated RES scores in the Riaz dataset. **d)** Expression levels of predictive genes that contain down-regulated RES scores in the Riaz dataset. *p<0.05, **p<0.01, ***p<0.001

**Figure S2:**
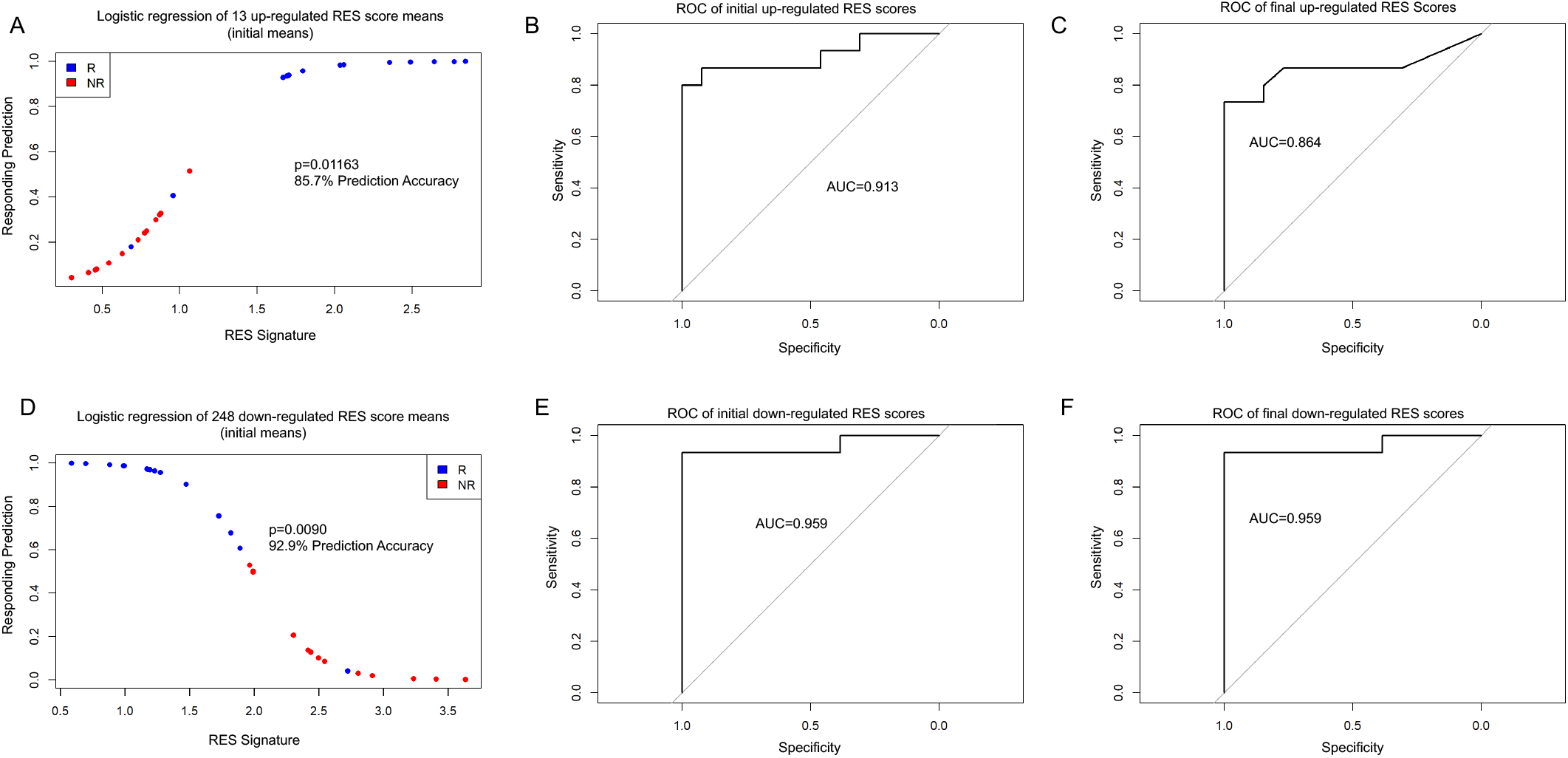
Logistic models and ROC curves from the Hugo dataset accurately predict patient response. **a)** Logistic regression models from the Hugo dataset for initial up-regulated RES score means and responding prediction for responder (R, blue) and non-responder (NR, red) patients. **b)** ROC curves of initial up-regulated RES score means. **c)** ROC curves of final up-regulated RES score means. **d)** Logistic regression models from the Hugo dataset for initial down-regulated RES score means and responding prediction for responder (R, blue) and non-responder (NR, red) patients. **e)** ROC curves of initial down-regulated RES score means. **f)** ROC curves of final down-regulated RES score means.

**Figure S3:**
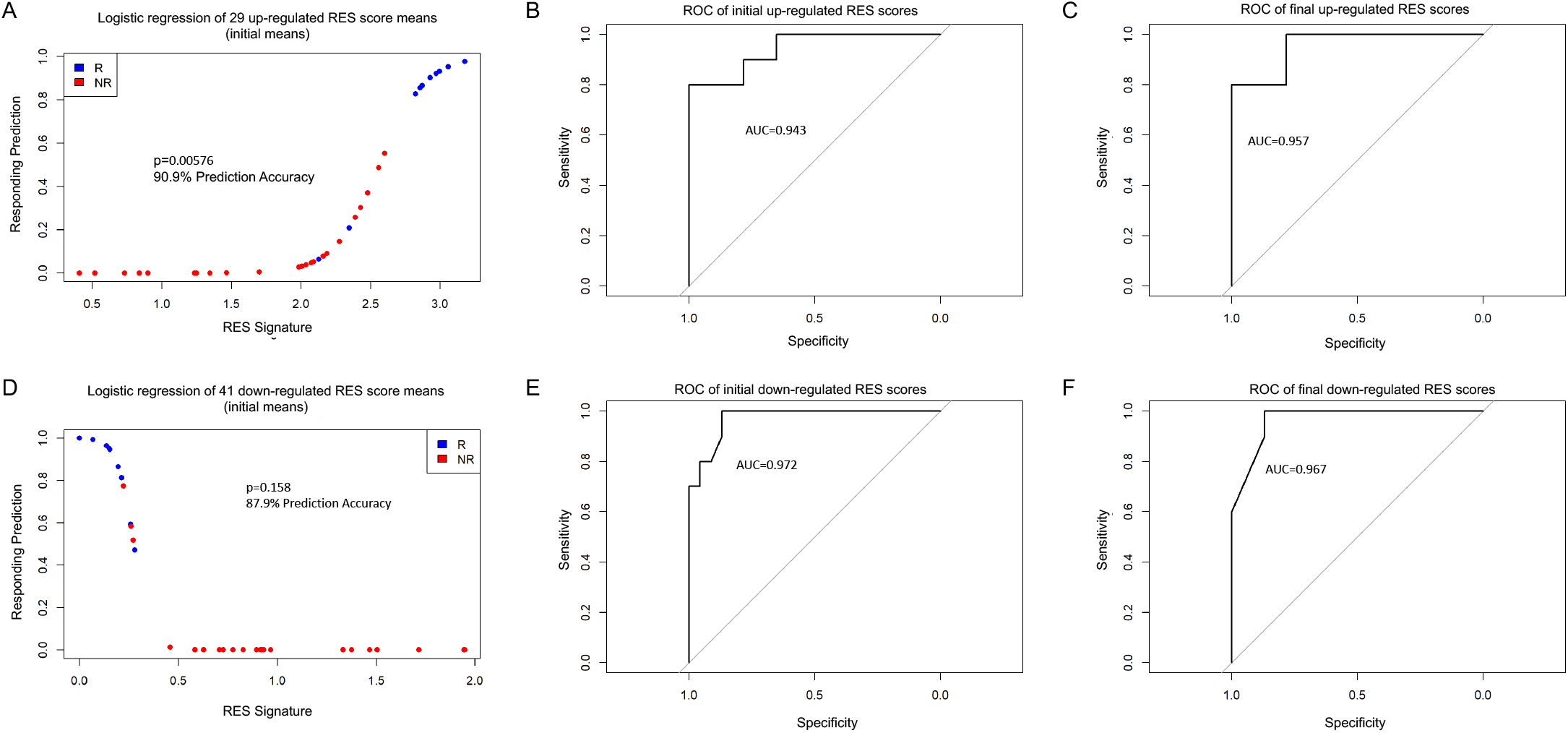
Logistic models and ROC curves from the Riaz dataset accurately predict patient response. **a)** Logistic regression models from the Riaz dataset for initial up-regulated RES score means and responding prediction for responder (R, blue) and non-responder (NR, red) patients. **b)** ROC curves of initial up-regulated RES score means. **c)** ROC curves of final up-regulated RES score means. **d)** Logistic regression models from the Riaz dataset for initial down-regulated RES score means and responding prediction for responder (R, blue) and non-responder (NR, red) patients. **e)** ROC curves of initial down-regulated RES score means. **f)** ROC curves of final down-regulated RES score means.

**Figure S4:**
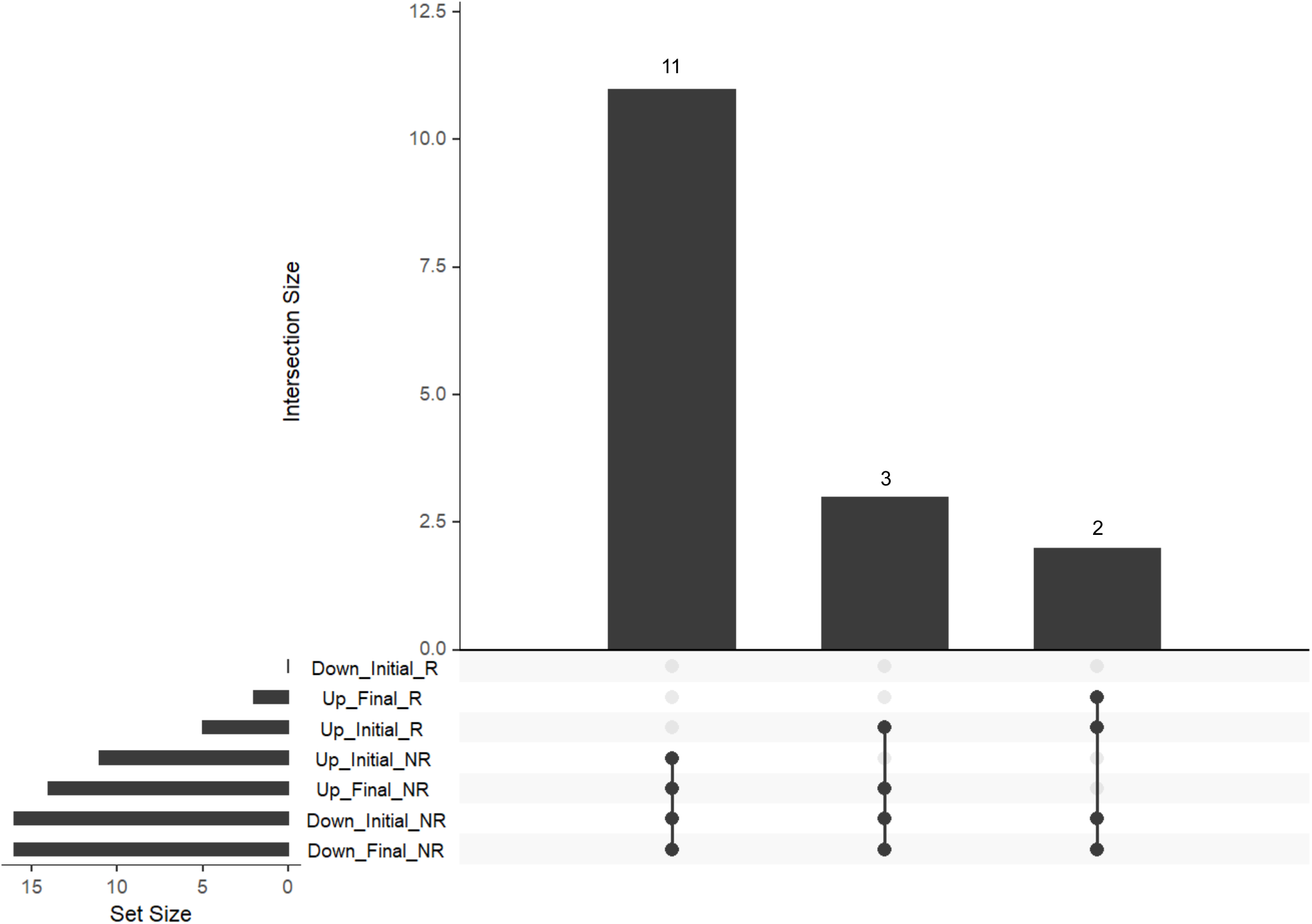
Logistic regression models cluster patients with stable disease with non-responders. Graph showing the frequency of stable disease RES scores clustering with non-responders (NR) and responders (R).

**Table S1:**
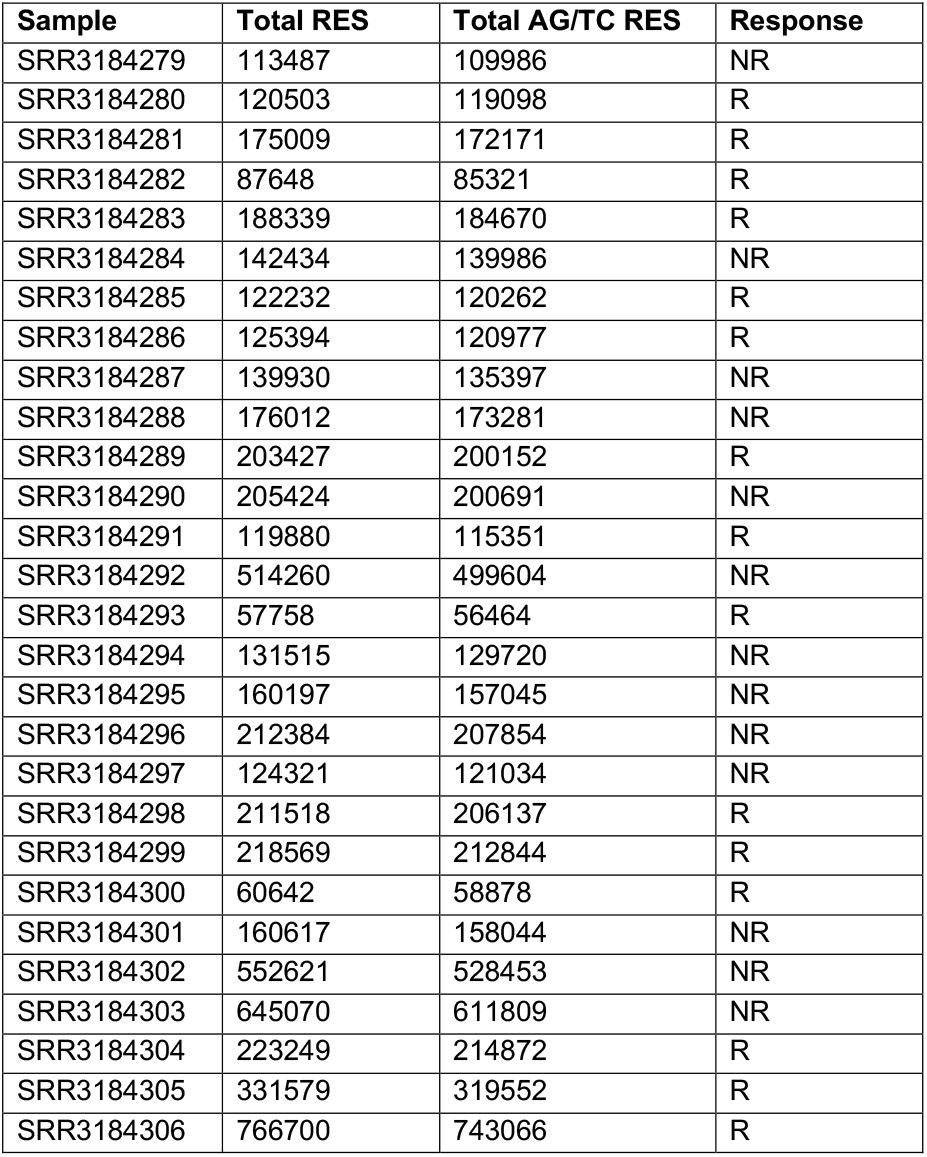
List of Hugo et al, 2016 samples and total RES identified

| Sample | Total RES | Total AG/TC RES | Response |
| --- | --- | --- | --- |
| SRR3184279 | 113487 | 109986 | NR |
| SRR3184280 | 120503 | 119098 | R |
| SRR3184281 | 175009 | 172171 | R |
| SRR3184282 | 87648 | 85321 | R |
| SRR3184283 | 188339 | 184670 | R |
| SRR3184284 | 142434 | 139986 | NR |
| SRR3184285 | 122232 | 120262 | R |
| SRR3184286 | 125394 | 120977 | R |
| SRR3184287 | 139930 | 135397 | NR |
| SRR3184288 | 176012 | 173281 | NR |
| SRR3184289 | 203427 | 200152 | R |
| SRR3184290 | 205424 | 200691 | NR |
| SRR3184291 | 119880 | 115351 | R |
| SRR3184292 | 514260 | 499604 | NR |
| SRR3184293 | 57758 | 56464 | R |
| SRR3184294 | 131515 | 129720 | NR |
| SRR3184295 | 160197 | 157045 | NR |
| SRR3184296 | 212384 | 207854 | NR |
| SRR3184297 | 124321 | 121034 | NR |
| SRR3184298 | 211518 | 206137 | R |
| SRR3184299 | 218569 | 212844 | R |
| SRR3184300 | 60642 | 58878 | R |
| SRR3184301 | 160617 | 158044 | NR |
| SRR3184302 | 552621 | 528453 | NR |
| SRR3184303 | 645070 | 611809 | NR |
| SRR3184304 | 223249 | 214872 | R |
| SRR3184305 | 331579 | 319552 | R |
| SRR3184306 | 766700 | 743066 | R |

**Table S2:**
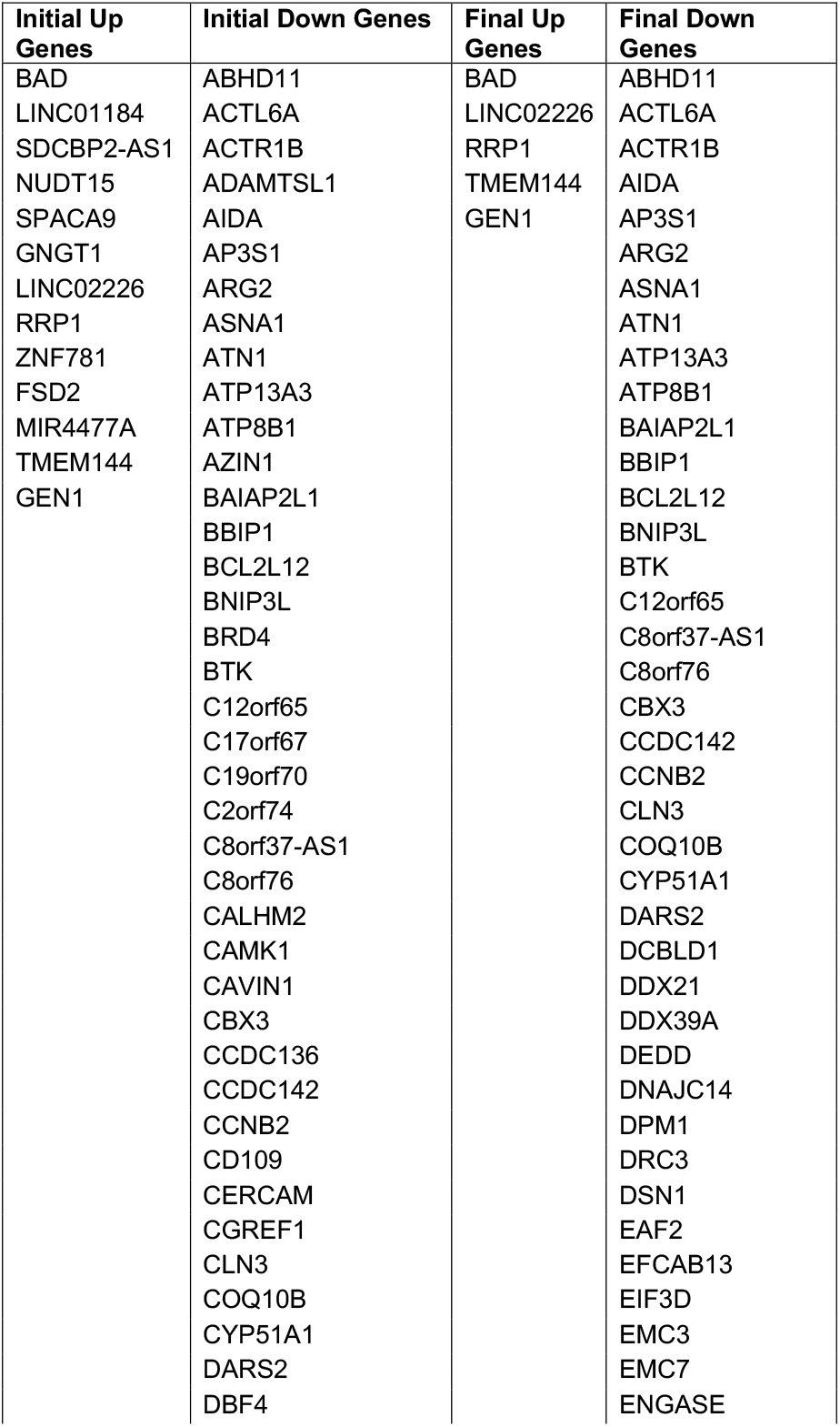
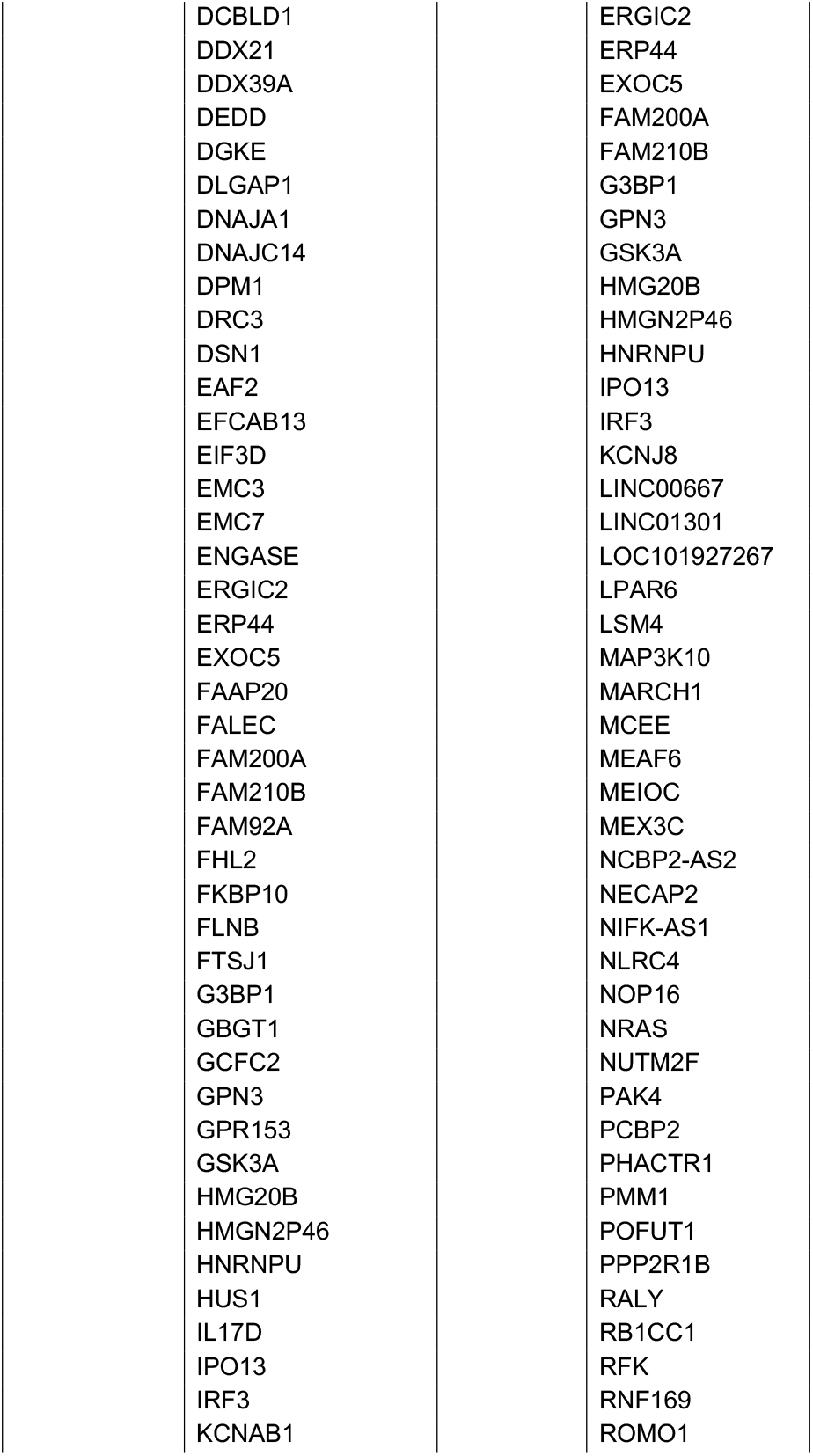
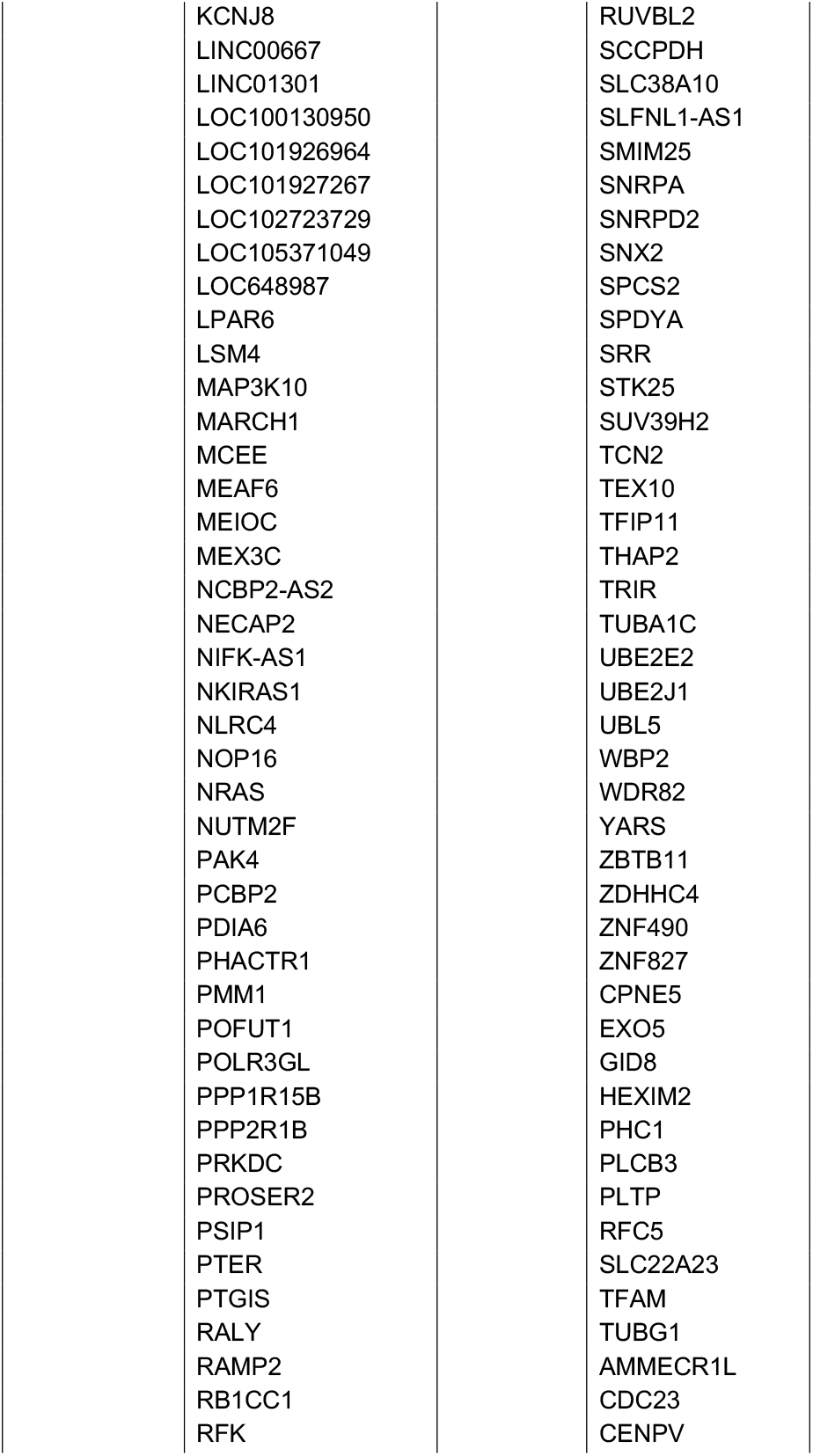
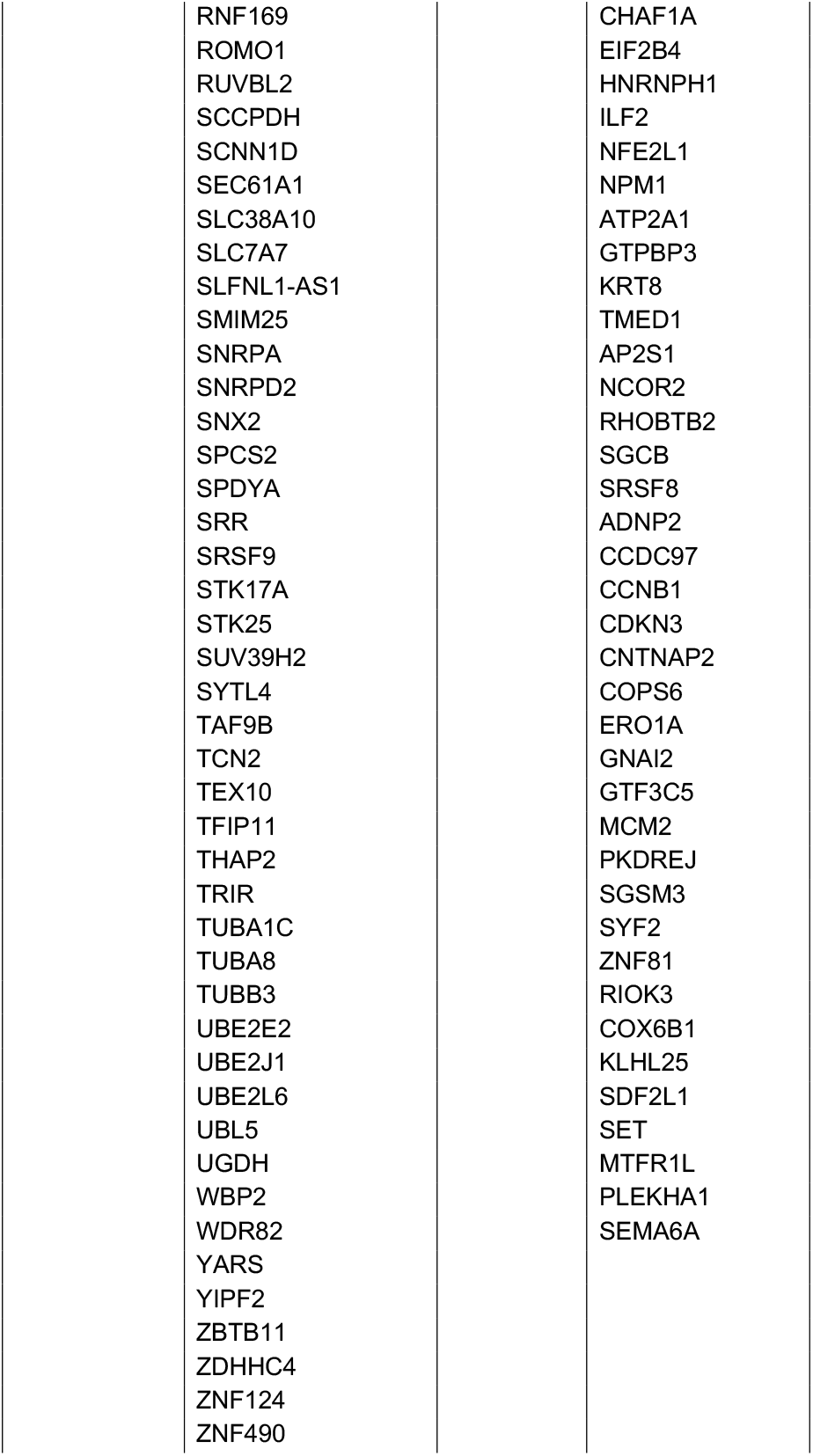
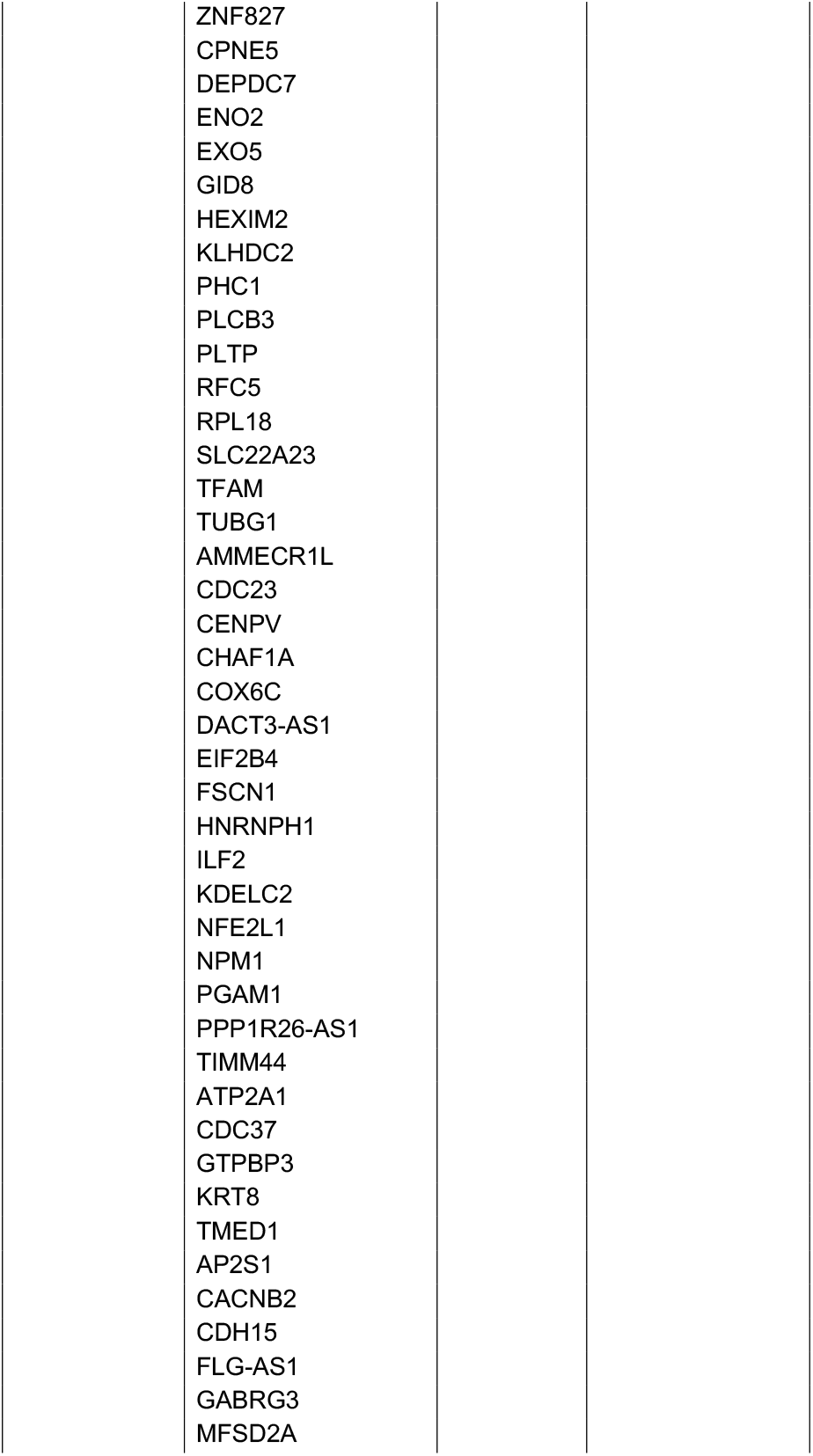
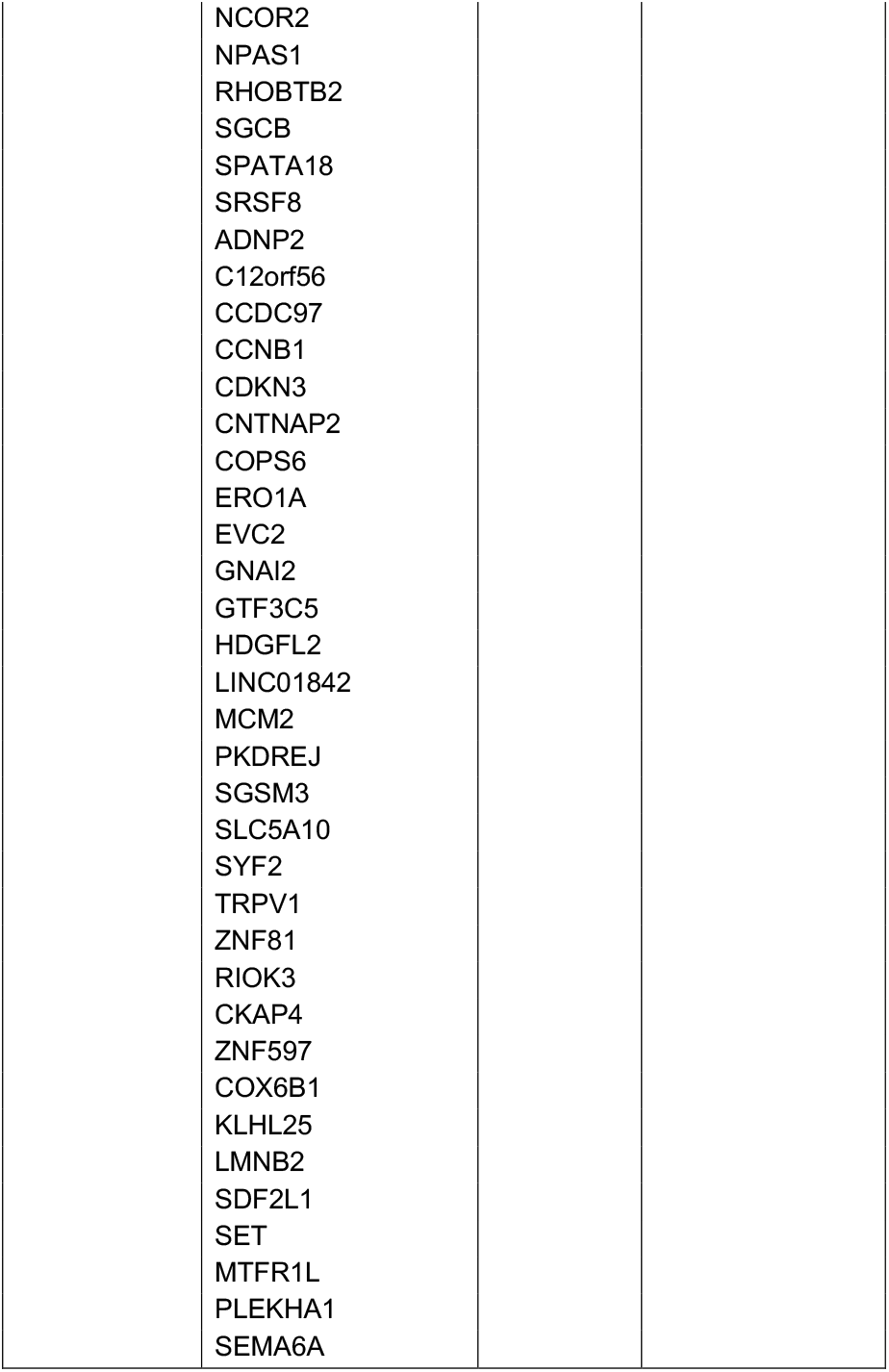
RES score signature genes from Hugo et al, 2016 cohort

**Table S3:** T-test of Hugo et al 2016 model predictions

| Model | Group0 | Group1 | D.O.F. | Mean Group0 | Mean Group1 | T-test Statistic | T-test P-value | Diff. of Means |
| --- | --- | --- | --- | --- | --- | --- | --- | --- |
| Initial Up RES Scores | NR | R | 17.02 | 0.67 | 1.86 | 6.00 | 1.44E-05 | 1.19 |
| Initial Down RES Scores | NR | R | 25.43 | 2.63 | 1.32 | -6.33 | 1.17E-06 | -1.30 |
| Final Up RES Scores | NR | R | 19.13 | 0.39 | 1.41 | 4.44 | 2.76E-04 | 1.02 |
| Final Down RES Scores | NR | R | 25.29 | 2.52 | 1.22 | -6.34 | 1.18E-06 | -1.30 |
| Final Up Transcript Levels | NR | R | 25.58 | 7.56 | 7.72 | 1.50 | 1.46E-01 | 0.16 |
| Final Down Transcript Levels | NR | R | 25.98 | 10.07 | 10.02 | -1.69 | 1.02E-01 | -0.05 |

D.O.F. (Degrees of Freedom)

**Table S4:** List of Riaz et al, 2017 samples and total RES identified

| Sample | Total RES | Total AG/TC Sites | Response |
| --- | --- | --- | --- |
| SRR5088813 | 14560 | 14119 | NR |
| SRR5088887 | 29306 | 28448 | SD |
| SRR5088834 | 9641 | 9185 | R |
| SRR5088872 | 1812 | 1678 | NR |
| SRR5088926 | 22203 | 21496 | NR |
| SRR5088916 | 16108 | 15677 | NR |
| SRR5088890 | 21140 | 20613 | NR |
| SRR5088891 | 11083 | 10684 | R |
| SRR5088815 | 14753 | 14388 | SD |
| SRR5088818 | 9335 | 9032 | UNK |
| SRR5088906 | 9075 | 8765 | NR |
| SRR5088920 | 35971 | 35303 | SD |
| SRR5088908 | 3964 | 3723 | NR |
| SRR5088922 | 10632 | 10323 | NR |
| SRR5088909 | 12812 | 12475 | NR |
| SRR5088878 | 13230 | 12797 | R |
| SRR5088911 | 25997 | 25426 | R |
| SRR5088913 | 13864 | 13503 | NR |
| SRR5088929 | 9690 | 9396 | R |
| SRR5088898 | 10108 | 9806 | SD |
| SRR5088900 | 16295 | 15830 | SD |
| SRR5088895 | 21154 | 20687 | SD |
| SRR5088819 | 9931 | 9643 | NR |
| SRR5088880 | 18138 | 17665 | SD |
| SRR5088924 | 16604 | 16255 | R |
| SRR5088904 | 10706 | 10435 | NR |
| SRR5088836 | 22165 | 21643 | NR |
| SRR5088897 | 17623 | 17268 | R |
| SRR5088824 | 13629 | 13326 | R |
| SRR5088883 | 6286 | 6023 | NR |
| SRR5088839 | 8651 | 8351 | NR |
| SRR5088821 | 13001 | 12587 | SD |
| SRR5088856 | 4934 | 4644 | NR |
| SRR5088857 | 22499 | 21940 | SD |
| SRR5088826 | 22415 | 21708 | NR |
| SRR5088840 | 6187 | 6025 | SD |
| SRR5088861 | 8060 | 7798 | R |
| SRR5088827 | 13043 | 12788 | UNK |
| SRR5088846 | 7804 | 7475 | SD |
| SRR5088864 | 4821 | 4367 | NR |
| SRR5088849 | 22065 | 21597 | SD |
| SRR5088914 | 17364 | 16941 | NR |
| SRR5088822 | 7025 | 6704 | SD |
| SRR5088850 | 9718 | 9364 | NR |
| SRR5088829 | 16919 | 16451 | NR |
| SRR5088831 | 9502 | 8961 | SD |
| SRR5088885 | 17741 | 17178 | NR |
| SRR5088866 | 10496 | 10181 | NR |
| SRR5088867 | 19327 | 18877 | SD |
| SRR5088843 | 8668 | 8336 | R |
| SRR5088853 | 11952 | 11545 | SD |

**Table S5:** RES score signature Genes from Riaz et al, 2017 cohort

| Initial Up Genes | Initial Down Genes | Final Up Genes | Final Down Genes |
| --- | --- | --- | --- |
| ADGRE2 | ACYP1 | C16orf72 | CFAP36 |
| AUH | ADCY6 | EIF2S3 | COX7B |
| C16orf72 | C2CD3 | ATOX1 | RAD50 |
| CTSS | CFAP36 | GOLPH3L | UBR4 |
| DTWD2 | COX7B | FCGRT | WASHC5 |
| EIF2S3 | GIT2 | ACBD4 | OSGEPL1 |
| FAM129A | GNS | CHADL | WAC |
| FAM20B | ITCH | ZNF236-DT | RBM10 |
| FPGS | LINC02614 | NAA40 |  |
| H6PD | MLANA |  |  |
| KRIT1 | MORC3 |  |  |
| LSG1 | MYO5A |  |  |
| PINK1-AS | RAB17 |  |  |
| POLR1E | RAD50 |  |  |
| RALGAPA1 | SEC16B |  |  |
| SOD2 | THUMPD3-AS1 |  |  |
| TEP1 | TSPAN31 |  |  |
| TRIM56 | UBR4 |  |  |
| ATOX1 | WASHC5 |  |  |
| FAM111A-DT | ZNF226 |  |  |
| GOLPH3L | ZNF329 |  |  |
| IL12RB1 | ZNF426 |  |  |
| LRRC57 | ZNF836 |  |  |
| ST3GAL2 | GMEB1 |  |  |
| FCGRT | GXYLT2 |  |  |
| ACBD4 | OSGEPL1 |  |  |
| CHADL | PRPF3 |  |  |
| ZNF236-DT | SVIL-AS1 |  |  |
| NAA40 | WAC |  |  |
|  | ZNF814 |  |  |
|  | ZNF718 |  |  |
|  | AKAP10 |  |  |
|  | FMNL2 |  |  |
|  | GOLGA2 |  |  |
|  | LOC100130950 |  |  |
|  | BICD1 |  |  |
|  | ABCA5 |  |  |
|  | GON4L |  |  |
|  | GYG2 |  |  |
|  | RAF1 |  |  |
|  | RBM10 |  |  |

**Table S6:** T-test of Riaz et al 2017 model predictions

| Model | Group0 | Group1 | D.O.F. | Mean Group0 | Mean Group1 | T-test Statistic | T-test P-value | Mean of Diff. |
| --- | --- | --- | --- | --- | --- | --- | --- | --- |
| Initial Up RES Score | NR | R | 30.65 | 1.72 | 2.82 | 6.16 | 8.16E-07 | 1.09 |
| Initial Up RES Score | NR | SD | 36.99 | 1.72 | 2.24 | 2.72 | 9.84E-03 | 0.51 |
| Initial Up RES Score | SD | R | 23.77 | 2.24 | 2.82 | 3.66 | 1.26E-03 | 0.58 |
| Initial Down RES Score | NR | R | 25.02 | 0.99 | 0.17 | -7.42 | 9.01E-08 | -0.81 |
| Initial Down RES Score | NR | SD | 30.81 | 0.99 | 0.91 | -0.47 | 6.39E-01 | -0.08 |
| Initial Down RES Score | SD | R | 16.29 | 0.91 | 0.17 | -5.23 | 7.73E-05 | -0.73 |
| Final Up RES Score | NR | R | 20.38 | 0.87 | 2.03 | 7.27 | 4.38E-07 | 1.15 |
| Final Up RES Score | NR | SD | 35.59 | 0.87 | 1.19 | 2.29 | 2.84E-02 | 0.32 |
| Final Up RES Score | SD | R | 19.27 | 1.19 | 2.03 | 5.26 | 4.29E-05 | 0.83 |
| Final Down RES Score | NR | R | 28.00 | 1.04 | 0.13 | -7.14 | 8.96E-08 | -0.90 |
| Final Down RES Score | NR | SD | 26.68 | 1.04 | 0.81 | -1.05 | 3.05E-01 | -0.23 |
| Final Down RES Score | SD | R | 16.95 | 0.81 | 0.13 | -3.56 | 2.41E-03 | -0.67 |
| Final Up Transcript Levels | NR | R | 13.25 | 9.55 | 9.52 | -0.26 | 7.96E-01 | -0.03 |
| Final Up Transcript Levels | NR | SD | 33.87 | 9.55 | 9.63 | 1.00 | 3.23E-01 | 0.08 |
| Final Up Transcript Levels | SD | R | 14.11 | 9.63 | 9.52 | -0.89 | 3.89E-01 | -0.11 |
| Final Down Transcript Levels | NR | R | 17.63 | 10.73 | 10.72 | -0.20 | 8.43E-01 | -0.01 |
| Final Down Transcript Levels | NR | SD | 32.64 | 10.73 | 10.75 | 0.40 | 6.89E-01 | 0.02 |
| Final Down Transcript Levels | SD | R | 19.51 | 10.75 | 10.72 | -0.52 | 6.09E-01 | -0.04 |

D.O.F. (Degrees of Freedom)

**Table S7:** Logistic Regression Results

| **Cohort** | **Model** | **Estimate** | **Standard Error** | **Z value** | **P-value** | **Prediction** | **ROC AUC** |
| --- | --- | --- | --- | --- | --- | --- | --- |
| Hugo | Initial Up | 4.16 | 1.65 | 2.52 | 0.01 | 0.86 | 0.91 |
| Hugo | Initial Down | -4.30 | 1.65 | -2.61 | 0.01 | 0.93 | 0.96 |
| Hugo | Final Up | 2.76 | 1.04 | 2.65 | 0.01 | 0.79 | 0.86 |
| Hugo | Final Down | -4.46 | 1.70 | -2.62 | 0.01 | 0.96 | 0.96 |
| Riaz | Initial Up | 6.10 | 2.21 | 2.76 | 0.01 | 0.91 | 0.94 |
| Riaz | Initial Down | -23.66 | 16.75 | -1.41 | 0.16 | 0.88 | 0.97 |
| Riaz | Final Up | 5.91 | 2.20 | 2.69 | 0.01 | 0.91 | 0.96 |
| Riaz | Final Down | -12.02 | 5.86 | -2.05 | 0.04 | 0.88 | 0.97 |

**Table S8:** Cox Proportional Hazards Modeling Results

| Cohort | Model | Coefficient | exp(Coefficient) | Standard Error | Z Value | P-value |
| --- | --- | --- | --- | --- | --- | --- |
| Hugo | Final Up | -0.63 | 0.53 | 0.39 | -1.62 | 1.06E-01 |
| Hugo | Final Down | 0.92 | 2.50 | 0.35 | 2.60 | 9.35E-03 |
| Riaz | Final Up | -1.19 | 0.30 | 0.33 | -3.63 | 2.88E-04 |
| Riaz | Final Down | 0.75 | 2.11 | 0.24 | 3.09 | 2.00E-03 |
| Hugo | Common RES | -0.53 | 0.59 | 0.22 | -2.42 | 1.53E-02 |
| Riaz | Common RES | -0.63 | 0.53 | 0.26 | -2.38 | 1.73E-02 |

**Table S9:** Common RES Sites

| Chromosome | Start | End | Strand | Transition | Genes |
| --- | --- | --- | --- | --- | --- |
| chr1 | 155733924 | 155733925 | + | AG | DAP3 |
| chr2 | 151477764 | 151477765 | + | AG | RIF1 |
| chr2 | 24002029 | 24002030 | + | AG | UBXN2A |
| chr22 | 42383507 | 42383508 | - | TC | NFAM1 |
| chr7 | 92200451 | 92200452 | - | TC | KRIT1 |

